# On the limits to invasion prediction using coexistence outcomes

**DOI:** 10.1101/2023.03.23.533987

**Authors:** Jie Deng, Washington Taylor, Simon A. Levin, Serguei Saavedra

**Author notes:** **Competing financial interests** The authors declare no competing financial interests.

## Abstract

The dynamics of ecological communities in nature are typically characterized by probabilistic processes involving invasion dynamics. Because of technical challenges, however, the majority of theoretical and experimental studies have focused on coexistence dynamics. Therefore, it has become central to understand the extent to which coexistence outcomes can be used to predict analogous invasion outcomes relevant to systems in nature. Here, we study the limits to this predictability under a geometric and probabilistic Lotka-Volterra framework. We show that while survival probability in coexistence dynamics can be fairly closely translated into colonization probability in invasion dynamics, the translation is less precise between community persistence and community augmentation, and worse between exclusion probability and replacement probability. These results provide a guiding and testable theoretical framework regarding the translatability of outcomes between coexistence and invasion outcomes when communities are represented by Lotka-Volterra dynamics under environmental uncertainty.

## Introduction

The assembly of ecological communities depends on multiple factors associated with eco-evolutionary dynamics and randomness (Chang et al., 2021, Drake, 1991, Hubbell, 1997, Odum and Barrett, 2005, Schreiber and Rittenhouse, 2004, Song et al., 2021*a*, Vellend, 2016, Warren and Spencer, 1996). In nature, these factors are typically expressed under probabilistic invasion dynamics (i.e., adding new species across time under changing environmental conditions) (Deng et al., 2021, Fukami, 2015, Hill et al., 2004, Jones et al., 2022, Odum, 1969, Williamson and Fittera, 1996). Because of technical challenges, however, the majority of theoretical and experimental studies have focused on coexistence dynamics (i.e., having all species included at once under fixed environmental conditions) (Case, 2000, Clark et al., 2021, Friedman et al., 2017, Gause, 1932, MacArthur and Levins, 1967, Maynard et al., 2020, Vandermeer, 1975). For example, it has proved extremely difficult to maintain a stable initial community for a sufficient time to then do invasion experiments (Friedman et al., 2017, Gilpin, 1986, Gould et al., 2018, Park, 1954). Similarly, in the field, it becomes easier to record coexistence (or co-occurrence) outcomes rather than invasion outcomes (Barner et al., 2018). Therefore, it has become central to understand the extent to which the outcomes (e.g., individual survival, community persistence, competitor exclusion) of coexistence dynamics can be used to predict analogous outcomes (e.g., invader colonization, invader-resident augmentation, resident replacement) expected from invasion dynamics (Case, 1990, Fukami, 2015, Hubbell, 1997, Vagnon et al., 2022).

A key connection between coexistence and invasion dynamics is centered on the concept of *invasion growth rate* (Barabás et al., 2018, Case, 1990, Chesson and Kuang, 2008, Grainger and Gilbert, 2019, Law and Morton, 1996). The invasion growth rate quantifies the instantaneous rate at which an invader’s population will grow (positive values) or decline (negative values) right after its introduction into a community of resident species at equilibrium. The invasion growth rate assumes that an invader’s abundance (or density) is sufficiently small such that it does not affect the growth rate of resident species, but sufficiently large such that demographic stochasticity can be ignored (Barabás et al., 2018, Hofbauer and Schreiber, 2022). Therefore, positive invasion growth rates guarantee the successful short-term introduction of an invader species (Barabás et al., 2018). Under these assumptions and under pre-determined model parameterizations, coexistence outcomes can be derived from information about invasion outcomes and vice versa, answering whether an outcome can happen or not (Arnoldi et al., 2022, Hofbauer and Schreiber, 2022, Logofet, 1993, Serván and Allesina, 2021). While these criteria have provided key insights regarding the translatability between coexistence and invasion dynamics, these results can be difficult to validate in natural systems where outcomes and conditions are rarely the same (Angulo et al., 2021, Miller and Allesina, 2021, Song et al., 2021a, Zhao et al., 2021). This has revealed the importance of extending and developing a testable theory to test the translatability between invasion and coexistence dynamics under a probabilistic framework to answer how likely an outcome can happen (Chang et al., 2023, Deng et al., 2021, Jones et al., 2022, Williamson and Fittera, 1996).

In this paper, we study the limits to invasion prediction using coexistence outcomes, aiming to understand the extent to which the theories of invasion and coexistence are interchangeable. To this end, we develop a theoretical integration of invasion and coexistence under a geometric and probabilistic framework following generalized Lotka-Volterra dynamics. Recognizing the environmental uncertainties present in natural systems, our theoretical framework approaches invasion and coexistence dynamics from a probabilistic perspective. We illustrate an application of this integration by providing a theoretical expectation of the translatability between analogous probabilistic outcomes from coexistence and invasion dynamics. We show, geometrically and using an in silico experiment, that while individual survival probability (i.e., focal species survives) can be translated into invader colonization probability (i.e., invader species colonizes), there can be a large loss of translation between community persistence (i.e., focal species coexists with all non-focal species) and community augmentation (i.e., invader species coexists with all resident species), as well as between exclusion probability (i.e., focal species excludes any number of non-focal species) and replacement probability (i.e., invader species replaces any number of resident species). These results provide a guiding and testable theoretical framework regarding the (lack of) translatability between coexistence and invasion outcomes when communities are represented by Lotka-Volterra dynamics under environmental uncertainty.

### Theoretical framework

Our theoretical framework follows the central assumption that ecological dynamics can be described by the generalized Lotka-Volterra (gLV) model of the form

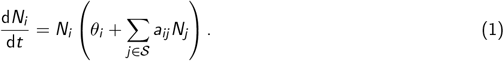

Here, 𝒮 is an ecological community (system) composed of |𝒮| species. The vector **N** = (*N*_1_, *…, N*_|𝒮|_)^T^ represents the densities of all species *i ∈ {*1, *…*, |𝒮|*}≡ 𝒮* in the community 𝒮. The matrix **A** = (*a*_*ij*_) *∈* R^|𝒮|*×*|𝒮|^ corresponds to the expected interaction matrix, where the entry *a*_*ij*_ represents the average percapita effect of species *j* on an individual of species *i*. That is, the interaction matrix **A** represents the expected biological structure within community 𝒮, consisting of the average interactions among species across time and space. In particular, *a*_*ij*_ *<* 0 and *a*_*ij*_ *>* 0 translate into a negative (competitive) and positive (cooperative) interaction, respectively. Thus, our theoretical framework can be applied to communities characterized by competition (where interactions *a*_*ij*_ *<* 0), cooperation (*a*_*ij*_ *>* 0), or a mix of both types of interactions. The effective growth rate ***θ*** = (*θ*_1_, *…, θ*_|𝒮|_)^*T*^ *∈* R^|𝒮|^ consists of the phenomenological joint effect of the internal (e.g., intrinsic growth rate), abiotic (e.g., temperature), and biotic factors (i.e., species not considered explicitly) acting on the per-capita growth rate of each particular species (Arnoldi et al., 2022, Deng et al., 2022).

Under our theoretical framework, we aim to address the challenges posed by the complexity in knowing and measuring the environmental factors affecting ecological communities. Instead of incorporating time-varying parameters or stochastic noise, we phenomenologically summarize the net effect of the environment on species using the vector of effective growth rates ***θ***. For a given community, the range of possible yet unknown environments can be mathematically mapped as the parameter space Θ^|𝒮|^ of effective growth rates ***θ*** for all the studied species. Without any prior information about the environments, we assume that all directions of ***θ*** are possible and equally likely, making the parameter space Θ^|𝒮|^ of ***θ*** a closed unit sphere in |𝒮|-dimensional space with a uniform probability distribution. Note that if we multiply ***θ*** by any positive scalar, it will not change the qualitative solution of the system. Thus, the choice of the norm (e.g., unit sphere) does not change the results (Rohr et al., 2016). Moreover, it is worth mentioning that our theoretical framework possesses the flexibility to be expanded to scenarios where environmental information is available. For instance, if the environments are known to support the growth of all species, then the parameter space Θ^|𝒮|^ can be constrained to the positive orthant. Alternatively, if the environments are distributed unevenly, with some being more likely to happen than others, one could use other geometric shapes such as an ellipsoid instead of a sphere to represent the parameter space Θ^|𝒮|^. More specifically, if the exact range of environments for a particular ecological community is known, one can in principle parameterize Θ^|𝒮|^ for coexistence and invasion dynamics separately based on this prior information.

In order to separate the role of environmental changes from community dynamics in shaping invasion and coexistence outcomes, we follow a structuralist (also known as internalist) perspective (Alberch, 1989, Kirschner and Gerhart, 2005). The structuralist perspective assumes that interactions between the individual components of a system are invariant and determine (or constrain) the environmental conditions compatible with the feasibility of living systems (Alberch, 1989). That is, while changes in a system are attributed to environmental (external) factors, the tolerance of a system to these changes is attributed to its biological (internal) structure (Alberch, 1991). In our ecological context, while it can be expected to observe changes in species interactions, the interaction matrix **A** can be interpreted as an expectation of an invariant internal structure of a community or complex living system (Rohr et al., 2014, Saavedra et al., 2017*b*).

In particular, in our gLV probabilistic framework, the uncertainty comes from the distribution of effective growth rates, but the interaction matrix determines the compatibility of a community with such uncertainty. We also restrict interaction matrices to cases of globally stable gLV dynamics (Takeuchi, 1996). Under this assumption, the results are independent of the initial conditions of the community and rely solely on the interaction matrix and effective growth rates (AlAdwani and Saavedra, 2020, Saavedra et al., 2020). This assumption further implies that any positive equilibrium of Eq. (1) guarantees the long-term persistence (also permanence) of species (Hofbauer and Sigmund, 1998). Notably, even in non-globally stable systems, our findings on the translatability between coexistence and invasion outcomes are qualitatively consistent (Supplementary Section S4). It is important to acknowledge that, while this assumption enables us to theoretically circumvent testing varying initial abundances, it is possible to investigate their effects on longterm outcomes in laboratory experiments (Gilpin, 1986). Additionally, a critical limitation to highlight is that our theoretical framework, under the assumption of global stability, does not accommodate the complex invasion phenomenon known as “residents strike back” (Mylius and Diekmann, 2001). Nevertheless, our proposed probabilistic frameworks has already been successfully applied to understand the behavior of communities that involve a wide range of interactions, dynamics, and populations (Allen-Perkins et al., 2023, Angulo et al., 2021, Bartomeus et al., 2021, Cenci et al., 2018, Deng et al., 2021, 2022, García-Callejas et al., 2023, Luo et al., 2022, Medeiros et al., 2021, Saavedra et al., 2017a, Song et al., 2018, 2017, Song and Saavedra, 2018, Song et al., 2023, 2021b, Tabi et al., 2020).

### Looking at invasion growth rate from a probabilistic perspective

To explore invasion dynamics, we borrow an existing measure known as the invasion growth rate that has been extensively studied and used in ecological theory, but we look at it from a probabilistic perspective. The invasion growth rate can accurately predict the behavior of an invader species over the short term. Under gLV dynamics Eq. (1), the invasion growth rate of an invader species *i* within a community 𝒮 formed by a pool of |𝒮| species can be formalized as

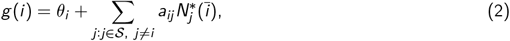

where 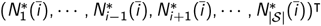 are the equilibrium abundances of the (|𝒮| *−* 1) resident species without the presence of invader species *i*. The definition of invasion growth rates implies that the invader species *i* has a low initial abundance. This assumption can be reasonable in nature as it is expected that at any given point in time only a small number of individuals can successfully migrate and reach the resident community. This definition also assumes the existence of steady-state positive abundances 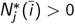 for all resident species *j ≠ i* in the absence of invader species *i*. Ecologically, the invasion growth rate (Eq. (2)) specifies the short-term population change of the invader species *i* when the resident species are at equilibrium. Assuming that resident species have reached equilibrium before the invasion event can offer valuable insights into the expected behavior of the resident community. This is because any changes that happen after the invasion can be attributed more confidently to the invader species, rather than the changing dynamics of the resident community. Per definition, the calculation of the invasion growth rate requires not only the interspecific effects of resident species on the invader species *a*_*ij*_, but also knowledge of the effective growth rate of the invader species *θ*_*i*_.

In the absence of prior information about *θ*_*i*_, we find that the invasion growth rate can be represented geometrically and probabilistically. To show this, first we define a geometric quantity, that we call the *growth indicator*, and then we associate this quantity with the invasion growth rate defined in Eq. (2). For an arbitrary vector of effective growth rates ***θ*** associated with the species forming a community 𝒮, the growth indicator of an invader species *i* can be defined as

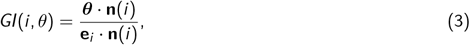

where **e**_*i*_ is the unit vector whose *i* ^th^ entry is 1 and 0 elsewhere; **n**(*i*) = (*n*_1_, *n*_2_, *…, n*_|𝒮|_)^*T*^ *∈* ℝ^|𝒮|^ can be any normal vector that is orthogonal to the *facet F* (*i*) in |𝒮|-dimensional space. Specifically, *F* (*i*) is spanned by the line segments from the origin to the points corresponding to the (|𝒮| *−* 1) column vectors excluding the *i* ^th^ column vector of the interaction matrix **A** (i.e., *F* (*i*) is spanned by the line segments from the origin to the points corresponding to the 1^st^, *…*, (*i −* 1)^th^, (*i* + 1)^th^, *…*, |𝒮|^th^ column vectors, so that the normal vector **n**(*i*) is specific to the invader species *i*). By definition, any normal vector **n**(*i*) that is orthogonal to the facet *F* (*i*) has zero inner products with the vectors on the facet *F* (*i*). In addition, for vectors on the same side of the facet *F* (*i*), their inner products with the normal vector **n**(*i*) to the facet *F* (*i*) have the same sign. The sign of *GI*(*i, θ*) reveals whether the vector of effective growth rates ***θ*** is on the same side of *F* (*i*) as the unit vector **e**_*i*_ relative to the facet *F* (*i*). It can be proved that *GI*(*i, θ*) is equivalent to the invasion growth rate *g* (*i*) (Supplementary Section S1). More specifically, if ***θ*** is on the same (resp. opposite) side as **e**_*i*_ (i.e., *GI*(*i, θ*) *>* 0; resp. *GI*(*i, θ*) *<* 0), then the invasion growth rate is *g* (*i*) *>* 0 (resp. *g* (*i*) *<* 0). Otherwise, if ***θ*** is on the facet *F* (*i*) (i.e., *GI*(*i, θ*) = 0), then the invasion growth rate is *g* (*i*) = 0. That is, the area spanned by *GI*(*i, θ*) *>* 0 corresponds to the range of conditions leading to a positive invasion growth rate.

Figure 1A illustrates geometrically the regions in the parameter space of ***θ*** corresponding to positive and negative invasion growth rates for a hypothetical community with three species. In this representation, we assume species 3 is the invader and species *{*1, 2*}*are the residents. The interaction matrices are arbitrary, and the plane *F* (*i*) where the invasion growth rate *g* (*i*) = 0 is determined by the 1^st^ and 2^nd^ column vectors corresponding to the resident species. Focusing on the feasibility domain of resident species in isolation (bounded by the green dotted lines), if the vector of effective growth rates ***θ*** is located inside the gray region *GI*(*i, θ*) *>* 0 (resp. white region *GI*(*i, θ*) *<* 0), the invasion growth rate is positive (resp. negative), meaning that the invader’s population with a small initial abundance will grow (resp. decline) in the short term. Otherwise, if the vector of effective growth rates ***θ*** is located at the border (i.e., on facet *F* (*i*)) dividing the gray and white regions, the invasion growth rate is zero, meaning that the invader’s population with a small initial abundance will remain unchanged over a short term. In the absence of prior information, we can assume that all values of ***θ*** are equally likely. Then, the probability that species *i* enters into a resident community with a positive invasion growth rate (*g* (*i*) *>* 0) can be measured by the area of ***θ*** compatible with *GI*(*i, θ*) *>* 0 (gray region) relative to the area of ***θ*** compatible with the persistence of all resident species (region bounded by the green dotted lines). Mathematically, in |𝒮|-dimensional space, the set of effective growth rates ***θ*** allowing the persistence of a resident community *𝒮 \ {i}*in isolation is

**Figure 1:**
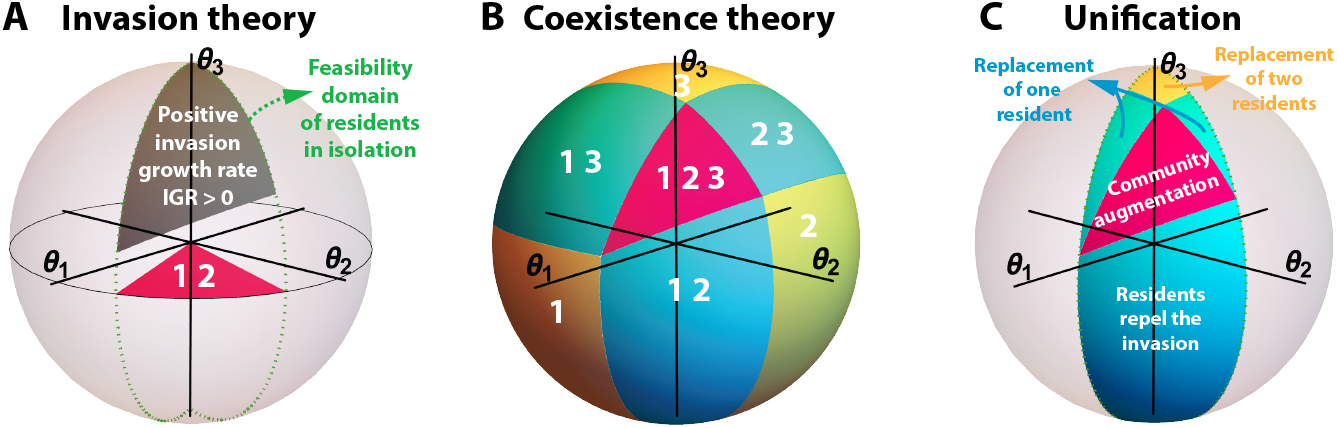
Geometric and probabilistic framework under generalized Lotka-Volterra dynamics. The figure illustrates our framework for a hypothetical three-species community. The sphere represents the parameter space of effective growth rates ***θ***, in which we color-code the regions compatible with feasible solutions 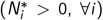 for different sets of species under the generalized Lotka-Volterra dynamics (Eq. (1)). In this illustration, resident species are labeled *{*1, 2*}*and the invader species*{* 3*}*. Panel A shows the feasibility domain (pink area) of 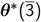 for resident species in the absence of the invader species. Note that this region is on the plane (*θ*_1_, *θ*_2_). The area spanned by the green dotted lines represents the region of the three-dimensional sphere that projects into this two-dimensional feasibility region. Then, the gray area corresponds to the region compatible with a positive invasion growth rate for the invader species (Eqs. (2-6)) assuming that the pre-existing resident species are at a feasible equilibrium (area constrained inside the green dotted lines). Panel B illustrates the hypothetical feasibility domains compatible with the long-term persistence of different collections of species (labeled on top of each region) assuming that all directions of ***θ*** are possible (Eqs. (7-8)). Panel C shows the possible long-term persistence outcomes under a successful or unsuccessful short-term invasion (combining previous panels): community augmentation, replacement of one or two resident species, and residents repelling the invasion (Eq. (9)). The possible outcomes are restricted to the area spanned by the feasibility of resident species in isolation.

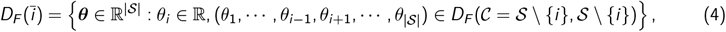

where the feasibility domain *D*_*F*_ (*𝒞, 𝒮*) of a community 𝒞 (*𝒞 ⊆ 𝒮, 𝒞 ≠ ∅*) within a multispecies community 𝒮 is defined in Eq. (7). Without prior information, we assume that the different directions of ***θ*** in 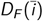 are equally likely, and consider these directions as a sub-region of the closed unit sphere in dimension |𝒮|.

Thus, the feasibility domain of resident species in isolation 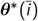 can be represented by

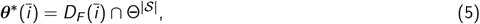

where Θ^|𝒮|^ is the (|𝒮| *−* 1)-dimensional closed unit sphere in dimension |𝒮|. For example, the region bounded by the green dotted lines in Fig. 1A is the feasibility domain of resident species *{*1, 2*}*in isolation, denoted by 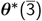. Formally, the probability of an invader species *i* having positive invasion growth rates in community |𝒮| under environmental uncertainty can be defined as

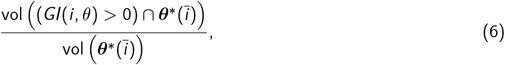

where “vol” means the volume of a region. In Fig. 1A, for example, the numerator in Eq. (6) corresponds to the volume of the gray region. Given the relationship between the growth indicator (*GI*(*i, θ*), Eq. (3)) and the invasion growth rate (*g* (*i*), Eq. (2)), this probability can be used to define the colonization probability of the invader species (Eq. (13)). Interestingly, our findings indicate that the probability of positive invasion growth rate (Eq. (6)) can be partitioned into multiple probabilities. Each of these corresponds to a specific outcome following the successful entry of the invader species. The partition is detailed in the following section.

### Looking at coexistence from a probabilistic perspective

As previously discussed, the invasion growth rate determines the invasion dynamics over the short term. Yet, the establishment of an invader species and its effects on resident species require the analysis of species persistence over a longer time scale. Under global equilibrium dynamics (Takeuchi, 1996), feasibility (positive solutions to Eq. (1)) constitutes the necessary and sufficient condition for species persistence (and permanence) (Hofbauer and Sigmund, 1998). Following gLV dynamics, the set of vectors ***θ*** leading to feasible solutions has been called the feasibility domain (Saavedra et al., 2020). Formally, the feasibility domain *D*_*F*_ (*𝒞, 𝒮*) of a (sub)community *𝒞* (*𝒞 ⊆ 𝒮, 𝒞 ≠ ∅*) within a multispecies community *𝒮* characterized by an interaction matrix **A** can be represented by (Deng et al., 2022)

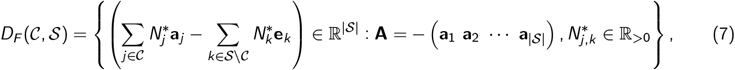

where **e**_*k*_ is a unit vector whose *k*^th^ entry is 1 and 0 elsewhere. Figure 1B illustrates geometrically the feasibility domains of different communities 𝒞 (which equivalently indicate the exclusion of species *k* not contained in 𝒞) within the same arbitrary community formed by three species as shown in Figure 1A. The column vectors of each interaction matrix determine the structure of the geometric partition (i.e., the feasibility domains). Specifically, when the vector of effective growth rates ***θ*** is located in the pink region labeled with 𝒞 = *{*1, 2, 3*}*, all three species are able to persist in the long term. Similarly, when the vector of effective growth rates ***θ*** is located in the blue region labeled with 𝒞 = *{*1, 2*}*, species 1 and 2 can persist and species 3 will be excluded in the long term.

Having no prior information about the values of ***θ*** (due to environmental uncertainty), we can again assume that all vector directions are equally likely and the feasibility domain *D*_*F*_ (*𝒞, 𝒮*) can be normalized by the volume of the entire parameter space (Saavedra et al., 2016). Then, under environmental uncertainty, the persistence probability over the long term of a community 𝒞 within a multispecies community 𝒮 can be analytically defined as (Deng et al., 2022)

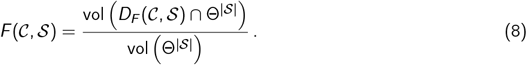

When 𝒞 = 𝒮, Eq. (8) corresponds to the persistence probability of the entire community 𝒮. Otherwise, when *𝒞 ⊂ 𝒮*, Eq. (8) corresponds to the persistence probability of a subcommunity 𝒞 only. If we consider community persistence as an outcome following probability theory, then the assigned probability (ranging from 0 to 1) for this outcome should be proportional to the sample space consisting of all possible outcomes. Here, the feasibility domain *D*_*F*_ (*𝒞, 𝒮*) corresponds to the persistence of a community 𝒞 within a larger community 𝒮. The sample space is the parameter space Θ^|𝒮|^ of effective growth rates ***θ*** in dimension |𝒮|. Therefore, the persistence probability of community 𝒞 can be represented by the volume (abbreviated as “vol” in Eq. (8)) of the feasibility domain *D*_*F*_ (*𝒞, 𝒮*) relative to the volume of the parameter space Θ^|𝒮|^. Ecologically, the persistence probability represents the fraction of compatible environmental contexts (***θ***) in which a community 𝒞 can persist itself within a larger community 𝒮, considering all possible environmental contexts.

### Integration of theories from a probabilistic perspective

To integrate invasion and coexistence theories, it is necessary to link short-term and long-term dynamics (Arnoldi et al., 2022, Hofbauer and Schreiber, 2022). From the invader’s perspective (species *i*), a positive invasion growth rate (i.e., *g* (*i*) *>* 0, measured as a function of the effect of resident species on the invader species, Eq. (2)) guarantees that its population will increase immediately after its introduction. For invader species *i*, the probability of a positive invasion growth rate is given by Eq. (6). However, the effect of a successful invader species on the resident species demands knowledge of how resident species will affect each other as well as how the invader species will affect the growth rate of resident species over the long run (Arnoldi et al., 2022, Hofbauer and Schreiber, 2022, Song et al., 2021a).

Following our geometric representation in the parameter space of ***θ*** (Fig. 1), we can illustrate the possible long-term effects of a successful invader species on the resident species by the colored regions in Figure 1C, for the same arbitrary community as shown in Figs. 1A and 1B. The pink region, where the entire community (i.e., resident species *{*1, 2*}*and invader species 3) can persist over the long term, corresponds to the domain of ***θ*** compatible with community augmentation. The two blue regions, where either resident 1 or resident 2 is excluded, correspond to the domain compatible with the replacement of one resident. The yellow region, where both resident species are excluded by the invader species, corresponds to the domain compatible with the replacement of two resident species. The region constrained by the green dotted lines corresponds to the environmental contexts compatible with the persistence of resident species in the absence of invader species *i* —the possible starting contexts for invasion.

Similar to the persistence probability (Eq. (8)) in coexistence theory, we can define probabilities to describe the chances of obtaining different outcomes within invasion dynamics under environmental uncertainty. Formally, after invader species *i* enters a resident community *𝒮 \{i}*, the probability of observing a community 𝒞 (*𝒞 ⊆ 𝒮*) can be defined as

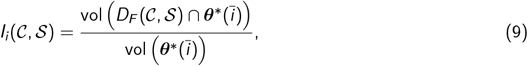

where 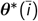 is the feasibility domain of resident species in isolation (without invader species *i*). We can understand Eq. (9) following the same idea as Eq. (8). Similarly, the feasibility domain *D*_*F*_ (*𝒞, 𝒮*) corresponds to the persistence of a community 𝒞 after the invasion of species *i*. However, the sample space now changes to the feasibility domain of resident species in isolation 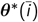, given that these species are present prior to any invasion event. Hence, the probability of observing community 𝒞 can be represented by the volume (abbreviated as “vol” in Eq. (9)) of the feasibility domain *D*_*F*_ (*𝒞, 𝒮*) relative to the volume of 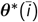. Ecologically, this probability represents the fraction of compatible environmental contexts (***θ***) where community 𝒞 can persist after the invasion of species *i*, considering all possible pre-invasion environmental contexts.

### Translating coexistence to invasion probabilistic outcomes

As an application of our integration of coexistence and invasion theories, we examine the extent to which we can use results from coexistence dynamics to infer results from invasion dynamics. For example, under coexistence dynamics, our theoretical framework can quantify the probabilities of survival of a focal species, community persistence, and the exclusion of non-focal species. Our theoretical framework can also describe analogous outcomes under invasion dynamics, which correspond to the probabilities of invader colonization, invader-resident augmentation, and resident replacement, respectively. For illustrative purposes, we focus on corresponding pairs of the aforementioned outcomes, but this study can be generalized to other outcomes.

Focusing on coexistence dynamics (Fig. 2, left column), the *probability of individual survival* of a focal species *i* (*i ∈ 𝒮*) can be calculated by the sum of feasibility domains (Eq. (7)) of all (sub)communities 𝒞 containing species *i* within the multispecies community 𝒮. Mathematically, this probability can be expressed as

**Figure 2:**
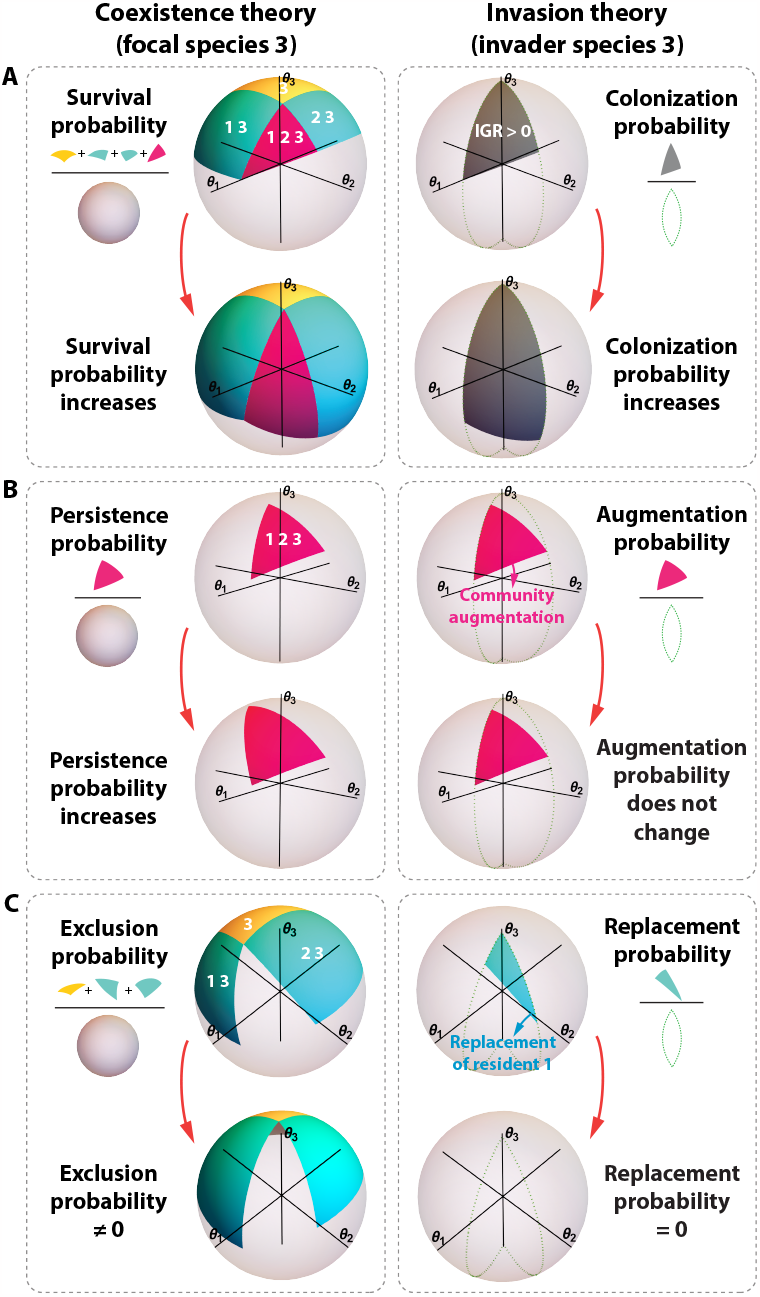
Geometric intuition on the translatability of analogous outcomes between coexistence and invasion dynamics. For a hypothetical three-species community, each panel (row) illustrates the relationship of analogous probabilistic outcomes between coexistence (left column, Eqs. (10-12)) and invasion (right column, Eqs. (13-15)) dynamics. The two spheres in each row correspond to the same community (i.e., the same interaction matrix **A**, Supplementary Section S2). Inside each panel, we represent an original and a changed community (i.e., a different **A**) at the top and bottom, respectively. Panel A shows the individual survival probability of focal species 3 (left, Eq. (10)) and the colonization probability of invader species 3 (right, Eq. (13)). In this example, both survival and colonization probabilities increase when resident species decrease their competitive effect on the invader species (change shown by red arrows). Panel B shows the persistence probability of the community (left, Eq. (11)) and the augmentation probability of the community (right, Eq. (14)). In this example, the persistence probability increases and the augmentation probability does not change when the competitive effects of the invader species on resident species decrease (change shown by red arrows). Panel C shows the exclusion probability that focal species 3 excludes non-focal species *{* 1,2*}*(left, Eq. (12)) and the replacement probability that invader species 3 replaces resident species *{* 1,2*}*(right, Eq. (15)). By decreasing the competitive effect of the invader species on resident species, this panel shows that the replacement probability is geometrically a small fraction of the exclusion probability (top) and can become zero when the exclusion probability is significantly greater than zero (bottom). Based on the geometric intuition illustrated here, there is an expected strong translatability between survival and colonization probabilities, but an expected weaker translatability between persistence and augmentation probabilities as well as between exclusion and replacement probabilities.

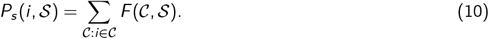

For example, considering a three-species community (𝒮 = *{*1, 2, 3*}*), the probability of survival of the focal species 3 is the summation of four feasibility domains: *P*_𝒮_ (3, *{*1, 2, 3*}*) = *F* (*{*3*}, {*1, 2, 3*}*)+*F* (*{*1, 3*}, {*1, 2, 3*}*)+ *F* (*{*2, 3*}, {*1, 2, 3*}*) + *F* (*{*1, 2, 3*}, {*1, 2, 3*}*) (Fig. 2A, left). Next, the *probability of community persistence* can be calculated by the feasibility domain where all species in the community 𝒮 can persist together.

Formally, it can be written as

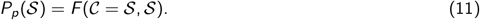

In this case, the probability of persistence of a three-species community (𝒮 = *{*1, 2, 3*}*) is given by just one feasibility domain: *P*_*p*_(*{*1, 2, 3*}*) = *F* (*{*1, 2, 3*}, {*1, 2, 3*}*) (Fig. 2B, left). Lastly, the *probability of exclusion* can be calculated by the difference between the probability of individual survival (Eq. (10)) and the probability of community persistence (Eq. (11), i.e., at least one non-focal species goes extinct, while the focal species survives). Mathematically, the probability of exclusion can be written as

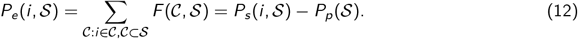

For example, considering a three-species community (𝒮 = *{*1, 2, 3*}*) with focal species 3, the probability of excluding at least one non-focal species is given by three feasibility domains: *P*_*e*_(3, *{*1, 2, 3*}*) = *F* (*{*3*}, {*1, 2, 3*}*) + *F* (*{*1, 3*}, {*1, 2, 3*}*) + *F* (*{*2, 3*}, {*1, 2, 3*}*) (Fig. 2C, left).

Shifting our focus to invasion dynamics (Fig. 2, right column), the *probability of invader colonization* of an invader species *i* can be calculated by the fraction of the feasibility domain of resident species 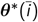 within which the invasion growth rate is positive. This probabilistic outcome is analogous to the probability of survival (Eq. (10)) under coexistence dynamics. Formally, it can be expressed as

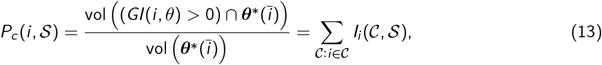

where the second relation holds under globally stable gLV dynamics (Eq. (1)). Results are qualitatively similar for locally stable communities (Supplementary Section S4). For example, considering species 3 as an invader and species *{*1, 2*}*as the resident community, the probability that the invader species can successfully colonize is given by *P*_*c*_ (3, *{*1, 2, 3*}*) = *I*_3_(*{*3*}, {*1, 2, 3*}*) + *I*_3_(*{*1, 3*}, {*1, 2, 3*}*) + *I*_3_(*{*2, 3*}, {*1, 2, 3*}*) +*I*_3_(*{*1, 2, 3*}, {*1, 2, 3*}*) (Fig. 2A, right). Next, the *probability of community augmentation* can be calculated by the fraction of the feasibility domain of resident species within which all species in the community 𝒮 can persist together and the invasion growth rate for species 3 is positive. This probabilistic outcome is analogous to the probability of community persistence (Eq. (11)) under coexistence dynamics. This probability can be written as

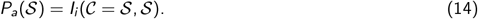

For example, the probability of community augmentation allowing invader species 3 to persist with the resident community *{*1, 2*}*is *P*_*a*_(*{*1, 2, 3*}*) = *I*_3_(*{*1, 2, 3*}, {*1, 2, 3*}*) (Fig. 2B, right). Lastly, the *probability of replacement* can be calculated by the sum of feasibility domains under which an invader species *i* can replace at least one resident species. This probabilistic outcome is analogous to the probability of exclusion (Eq. (12)) under coexistence dynamics. Mathematically, the probability that an invader species *i* replaces any resident is specified by

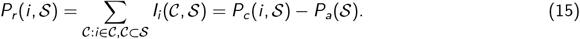

For instance, the probability that invader species 3 can replace at least one resident in the community *{*1, 2*}*is *P*_*r*_ (3, *{*1, 2, 3*}*) = *I*_3_(*{*3*}, {*1, 2, 3*}*) + *I*_3_(*{*1, 3*}, {*1, 2, 3*}*) + *I*_3_(*{*2, 3*}, {*1, 2, 3*}*) (Fig. 2C, right).

Importantly, our geometric framework can be used to gain insights about the extent to which probabilistic outcomes under coexistence dynamics can be used to obtain information regarding analogous outcomes under invasion dynamics. Figure 2 shows the geometric illustration of the three aforementioned pairs of analogous probabilities for hypothetical three-species communities (𝒮 = *{*1, 2, 3*}*). Similar geometric intuition can be applied to higher dimensions. The two spheres in each row correspond to the same three-species community 𝒮 (i.e., the same interaction matrix **A**, Supplementary Section S2), with specific but fairly arbitrary values of the interactions *a*_*ij*_, illustrating the possibility of translation between the relevant quantities. In each panel of Figure 2, we represent an original and changed community at the top and bottom, respectively—where in each case the change corresponds to a modification of certain entries of the interaction matrix **A** (Supplementary Section S2).

Figure 2A illustrates the pair formed by individual survival probability (Eq. (10)) and invader colonization probability (Eq. (13)). In this hypothetical case, an increase in the colonization probability is generally associated with an increase in the survival probability. Mathematically, the invasion growth rate (Eq. (2)) is an explicit function of the effects of resident species on the invader species. These effects are represented by the off-diagonal entries of the row vector corresponding to the invader species in the interaction matrix (i.e., the 3^rd^ row vector). For example, changing the effects of resident species on the invader species from competitive to cooperative (Fig. 2A, change shown by red arrows) moves the lower boundaries of the relevant regions downward in the lower spheres indicating the transformed system. This change consequently increases the region of positive invasion growth rates and is associated with an increase in both probabilities. This pair of analogous probabilities is also the simplest case compared to the other pairs, as both probabilities focus on one single species. Hence, we can expect a strong translatability between this pair of analogous probabilities.

Figure 2B shows the pair formed by community persistence probability (Eq. (11)) and community augmentation probability (Eq. (14)). In this second case, if the persistence probability increases, the augmentation probability does not necessarily increase. For example, changing the effects of the invader species on a resident species from competitive to cooperative (Fig. 2B, change shown by red arrows) increases the region of community persistence. But the region of positive invasion growth rates does not change. Also, the complexities involved in both persistence and augmentation probabilities are higher than the first pair of analogous probabilities, as they both focus on all three species in the community instead of one single species. This difference leads to an expected weaker translatability between this pair of analogous probabilities.

Figure 2C illustrates the pair formed by exclusion probability (Eq. (12)) and replacement probability (Eq. (15)). In this third case, the replacement probability can be significantly smaller than the exclusion probability. For example, changing the effects of the invader species on both resident species from competitive to cooperative (Fig. 2C, change shown by red arrows) decreases the exclusion probability, but can eliminate the possibility of replacement altogether. Furthermore, while this quantity is simply the difference of the other two quantities, the strongest correlative effect associated with changes in the effect of resident species on the invader species largely cancels. Additionally, the complexities involved in both exclusion and replacement probabilities are even higher due to the numerous outcomes based on the quantity and identity of species that become extinct. Taken together these effects suggest an expected weaker translatability between exclusion and replacement probabilities.

In sum, Figure 2 indicates a translatability between survival and colonization probabilities, but some loss of translation between persistence and augmented probabilities and even less correlation between exclusion and replacement probabilities. Note that our notion of translatability can be expressed in terms of predictability as correlations of computations of specific quantities given some known information about the system (e.g., this can be specified by the coefficient of determination, *R*^2^).

### In silico experiments

Next, we provide an *in silico* experiment as a way to illustrate the inputs and outputs required to corroborate our theoretical integration. Specifically, we perform numerical tests using simulations to confirm the loss of translatability between coexistence and invasion outcomes established by our geometric framework. To this end, we generate 100 *in silico* communities, each consisting of three species. This means generating 100 interaction matrices **A** of dimension 3 *×* 3, with interspecific effects (off-diagonal entries of **A**) being randomly selected from a normal distribution with mean *μ* = 0 and standard deviation *σ* = 0.5. The intraspecific effects (diagonal entries of **A**) are set to *a*_*ii*_ = *−*1. Thus, all three-species communities are globally stable under the gLV dynamics (Eq. (1)), making the results independent from the species’ initial abundances. These values are in line with the geometric illustration in Figure 2, but the qualitative results are robust to this choice (Supplementary Section S3).

All coexistence and invasion outcomes from these *in silico* communities can be calculated either analytically or numerically using their corresponding definitions (frequencies) established in Eqs. (10-15). The analytic calculation, however, is valid only in the idealized theoretical limit of an infinite number of experimental trials and time steps (Deng et al., 2022). Hence, we use a numerical approximation by simulations to make our analysis more comparable to the finite nature of lab/field experiments or observations (Deng et al., 2022). For this purpose, we randomly and uniformly sample 10^4^ vectors of effective growth rates ***θ*** for each community generated at random. Depending on the dynamics, these vectors are positioned either on the unit sphere Θ^|𝒮|^ for coexistence outcomes or within the feasibility domain of resident species in isolation 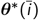 for invasion outcomes. Each one of these vectors mimics an experimental trial initiated under different environmental conditions. Under global stability, species with a positive equilibrium abundance can be classified as persistent (Takeuchi, 1996). This classification can also be reached via simulations after a given period of time assuming a given threshold for extinction (Deng et al., 2022). We simulate the gLV model (Eq. (1)) with total time 200, step size 10^*−*2^, and an extinction threshold of 10^*−*6^. This threshold may also correspond to the resolution level at which species can be detected in the lab or in the field (Deng et al., 2022). Simulations are conducted using the Runge-Kutta method. Hence, the probability of observing a community 𝒞 within a multispecies community 𝒮 (characterized by **A**), i.e., *F* (*𝒞, 𝒮*) in coexistence dynamics (Eq. (8)) and *I*_*i*_ (*𝒞, 𝒮*) in invasion dynamics (Eq. (9)), can be approximated by the frequency with which all species in the community 𝒞 persist together. In other words, this frequency is the fraction of successful trials out of all trials. Finally, using definitions shown in Eqs. (10-15), we can calculate the three pairs of analogous probabilities based on *F* (*𝒞, 𝒮*) and *I*_*i*_ (*𝒞, 𝒮*).

We use the coefficient of determination *R*^2^ to measure the translatability of analogous outcomes between coexistence and invasion dynamics. Panels A-C in Fig. 3 show, respectively, the relationship between individual survival probability and invader colonization probability (*R*^2^ = 0.98), community persistence probability and community augmentation probability (*R*^2^ = 0.74), and exclusion probability and replacement probability (*R*^2^ = 0.58). Importantly, this figure confirms the translatability between analogous outcomes expected from the geometric analysis (Fig. 2). That is, if one aims to use the outcomes from coexistence dynamics to predict the outcomes of invasion dynamics (or vice versa), the only highly reliable case would be the translation between survival and colonization probabilities.

**Figure 3:**
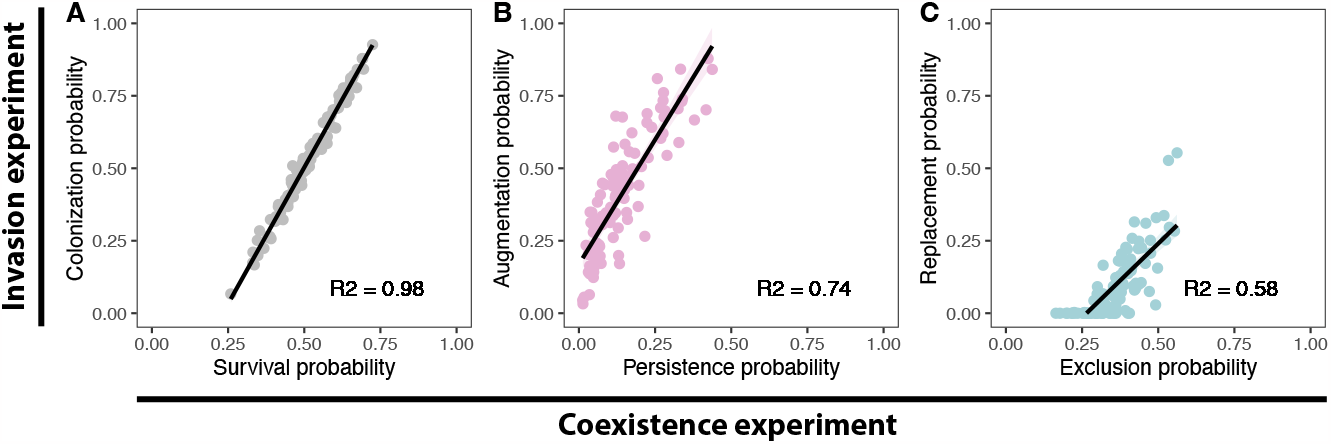
*In silico* experiment corroborating the (lack of) translatability between coexistence and invasion outcomes. The figure illustrates the numerical results using gLV simulations of three pairs of analogous probabilistic outcomes based on 100 randomly generated three-species communities (see main text for details). The x-axes and y-axes correspond to the outcomes under coexistence dynamics (Eqs. (10-12)) and invasion (Eqs. (13-15)) dynamics, respectively. Each point corresponds to a three-species community, whose interaction matrix **A** is randomly generated using a normal distribution with mean *μ* = 0 and standard deviation *σ* = 0.5 (diagonal entries *a*_*ii*_ =*−* 1). We also tested other parameters (e.g., community sizes, standard deviations) and obtained qualitatively the same results (Supplementary Section S3). These numerical tests confirm the expected strong translatability between survival and colonization probabilities (Panel A: *R*^2^ = 0.98), and the expected weaker translatability between persistence and augmentation probabilities (Panel B: *R*^2^ = 0.74) as well as between exclusion and replacement probabilities (Panel C: *R*^2^ = 0.58).

## Conclusions

Linking coexistence and invasion dynamics has been a fundamental scientific problem, underpinning our ability to predict the rise and fall of populations forming part of natural ecological communities (Fukami, 2015, Hastings, 2004, Odum, 1969, Vellend, 2016). Recent efforts to integrate these two types of dynamics have relied on the assumption that it is possible to fully parameterize ecological models (Arnoldi et al., 2022, Hofbauer and Schreiber, 2022), the assumption that environmental conditions remain the same across the entire process (Arnoldi et al., 2022, Hofbauer and Schreiber, 2022, Logofet, 1993, Serván and Allesina, 2021), or the assumption that one has sufficient information about coexistence or invasion outcomes (Friedman et al., 2017, Grainger and Gilbert, 2019, Maynard et al., 2020). While these studies have provided key insights regarding the association of coexistence and invasion dynamics, these assumptions are seldom met in nature (Fukami, 2015, Hubbell, 1997, Miller and Allesina, 2021, Song et al., 2021a). Therefore, it has become central to understand the extent to which the probabilistic outcomes from such coexistence dynamics are representative of analogous outcomes expected from more complex invasion dynamics. This translation, however, requires the non-trivial integration of coexistence and invasion theories under a probabilistic perspective.

To study the limits to invasion prediction using coexistence outcomes, we have integrated coexistence and invasion outcomes using a geometric and probabilistic framework (Fig. 1). We have derived this theoretical framework from generalized Lotka-Volterra dynamics assuming that effective per-capita growth rates are a function of the environment, considering a one-to-one mapping between the environment and effective per-capita growth rates, considering no a priori information about the environment (i.e., environmental uncertainty), and assuming that species interactions constrain the set of effective per-capita growth rates compatible with coexistence and invasion outcomes (following a structuralist perspective). As an application, we have used this framework to study the translatability from coexistence outcomes into invasion outcomes. We have provided probabilistic measures (Fig. 2) of coexistence outcomes (i.e., individual survival, community persistence, competitor exclusion) and invasion outcomes (i.e., invader colonization, invader-resident augmentation, resident replacement). We have shown that the translatability between the probability of analogous outcomes can be valid between survival and colonization. However, this translatability becomes weaker between persistence and augmentation, as well as between exclusion and replacement (Fig. 3). This suggests that outcomes from invasion dynamics involving more than one species should be treated differently from their coexistence counterparts. These results further support empirical observations showing a clear separation between coexistence and invasion dynamics (Angulo et al., 2021, Deng et al., 2021). Importantly, our integration provides a guiding and testable theoretical framework regarding the (lack of) translatability of outcomes between coexistence and invasion dynamics—which is relevant for a better understanding of systems in nature.

## S1 Growth indicator

Writing the generalized Lotka-Volterra model (Eq. (1)) in the matrix form 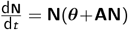, the interaction matrix **A** is an |𝒮| *×* |𝒮| matrix, i.e.,

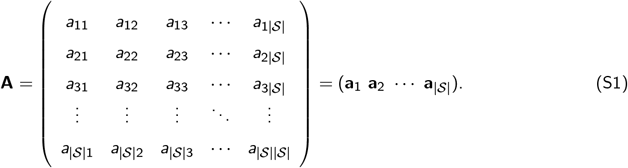

We assume, without loss of generality, species 1 as the invader and the other (|𝒮| *−* 1) species *{*2, *…*, |𝒮|*}*as the residents. Let **n** denote one of the normal vectors that is orthogonal to the facet *F* (*i*) (also called hyperface) spanned by the negative of the 2^nd^ to |𝒮|^th^ column vectors of the interaction matrix **A** (i.e., *−***a**_2_, *−***a**_3_, *…, −***a**_|𝒮|_). Note that the facet and its normal vector in the main text are denoted as *F* (*i*) and **n**(*i*), respectively (Eq. (3)) because the facet *F* (*i*) and its normal vector **n**(*i*) can change with invader species *i*. Here, for the convenience of reading, we simplify the facet and normal vector corresponding to invader species 1 as *F* and **n**. Specifically, any normal vector **n** that is orthogonal to the facet *F* has zero inner products with the vectors on the facet *F*. In addition, for vectors on the same side of the facet *F*, their inner products with the normal vector **n** to the facet *F* have the same sign. Moreover, to represent the facet *F*, we use the negative of the column vectors of the interaction matrix **A** which are also the spanning vectors of the feasibility domain *D*_*F*_ (*𝒞, 𝒮*) for easier geometric reference (see Eq. (7)).

Thus by definition, the following inner products are zero

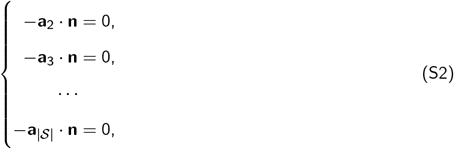

which is equivalent to the form removing the negative sign

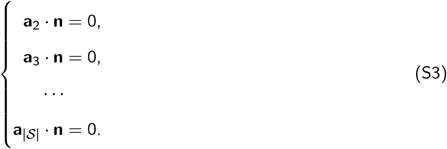

Let us denote **n** = (*n*_1_, *n*_2_, *…, n*_|𝒮|_)^*T*^, then the matrix form of Eq. (S3) can be written as

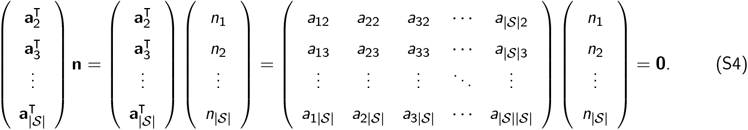

Next, we are going to simplify the matrix form Eq. (S4). First, let us set **a**_*inv*_ = (*a*_12_, *a*_13_, *…, a*_1|𝒮|_), 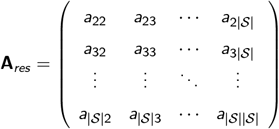, **n**_*res*_ = (*n*_2_, …, *n*_|S|_)^*T*^, then Eq. (S4) can be written as

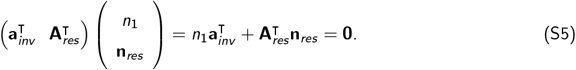

Hence,

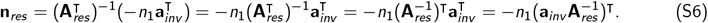

For an arbitrary effective growth rate ***θ***, we define the *growth indicator* as

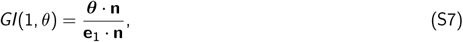

where **e**_1_ = (1, 0, …, 0)^*T*^. Apparently, a positive *GI*(1, *θ*) means that the inner products of the effective growth rate ***θ*** and the unit vector **e**_1_ with the normal vector **n** to the facet *F* have the same sign. This can be geometrically interpreted as the effective growth rate ***θ*** and the unit vector **e**_1_ are on the same side of the facet *F*.

Obviously, **e**_1_ *·* **n** = *n*_1_. Thus the growth indicator can be written as

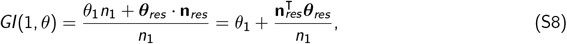

where ***θ***_*res*_ = (*θ*_2_, *…, θ*_|𝒮|_)^*T*^.

Substitute **n**_*res*_ by Eq. (S6), then

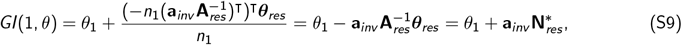

where 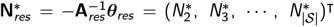 is the interior equilibrium of the resident community *{*2, 3, *…*, |𝒮|*}*in isolation. Note that the right side of Eq. (S9) is exactly the invasion growth rate *g* (1). Hence we prove that the growth indicator *GI*(1, *θ*) is equivalent to the invasion growth rate *g* (1). Since this proof also applies to a general invader species *i* in any dimension, it is safe to conclude that the growth indicator *GI*(*i, θ*) is equivalent to the invasion growth rate *g* (*i*). Therefore, the geometric representation of the region with positive invasion growth rate (i.e., *g* (*i*) *>* 0) is on the same side as the unit vector **e**_*i*_ (whose *i* ^th^ entry is 1 and 0 elsewhere) relative to the facet *F* (*i*) spanned by the (|𝒮| *−* 1) column vectors (except for the *i* ^th^ column vector) of the interaction matrix **A**.

## S2 Interaction matrices in Figure 2

As a reference, below are the interaction matrices **A** used in Figure 2, for the hypothetical communities consisting of three species. Further details are available in the main text.

- Figure 2A, before the change shown by red arrows 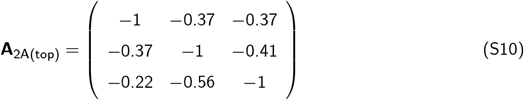
- Figure 2A, after the change shown by red arrows 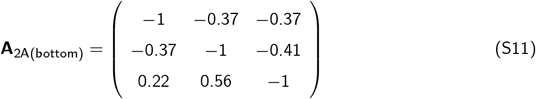
- Figure 2B, before the change shown by red arrows 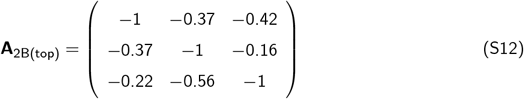
- Figure 2B, after the change shown by red arrows
- 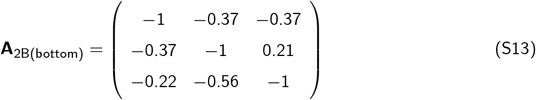
- Figure 2C, before the change shown by red arrows 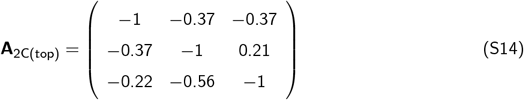
- Figure 2C, after the change shown by red arrows

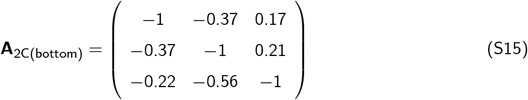

## S3 Robustness of results

In this section, we reproduce Fig. 3 with different numbers of species (dim) and different standard deviations of the interactions (sd). The figures illustrate the numerical results of three pairs of analogous probabilistic outcomes using 100 randomly generated communities. Specifically, each point corresponds to a community with the number of species (dim) *∈ {*3, 4, 5, 8, 10*}*. The corresponding interaction matrices **A** are randomly generated using a normal distribution with mean *μ* = 0 and standard deviations (sd) *∈ {*0.1, 0.5, 1, 1.5*}*(diagonal entries *a*_*ii*_ = *−*1). Specific parameter combinations are shown in the caption of each figure. Probabilities are calculated numerically using an ensemble of 10^4^ effective growth rates ***θ*** randomly sampled and then simulating the gLV dynamics (see main text for details). Then, the probabilities can be approximated by the frequencies that certain communities persist together over a finite period of time and threshold for species extinction (Deng et al., 2022). Consistent with the main text, we use the *R*^2^ to measure the translatability of analogous outcomes between coexistence and invasion dynamics. With different combinations of parameters, the numerical results are qualitatively the same as Fig. 3.

**Supplementary Figure S1:**
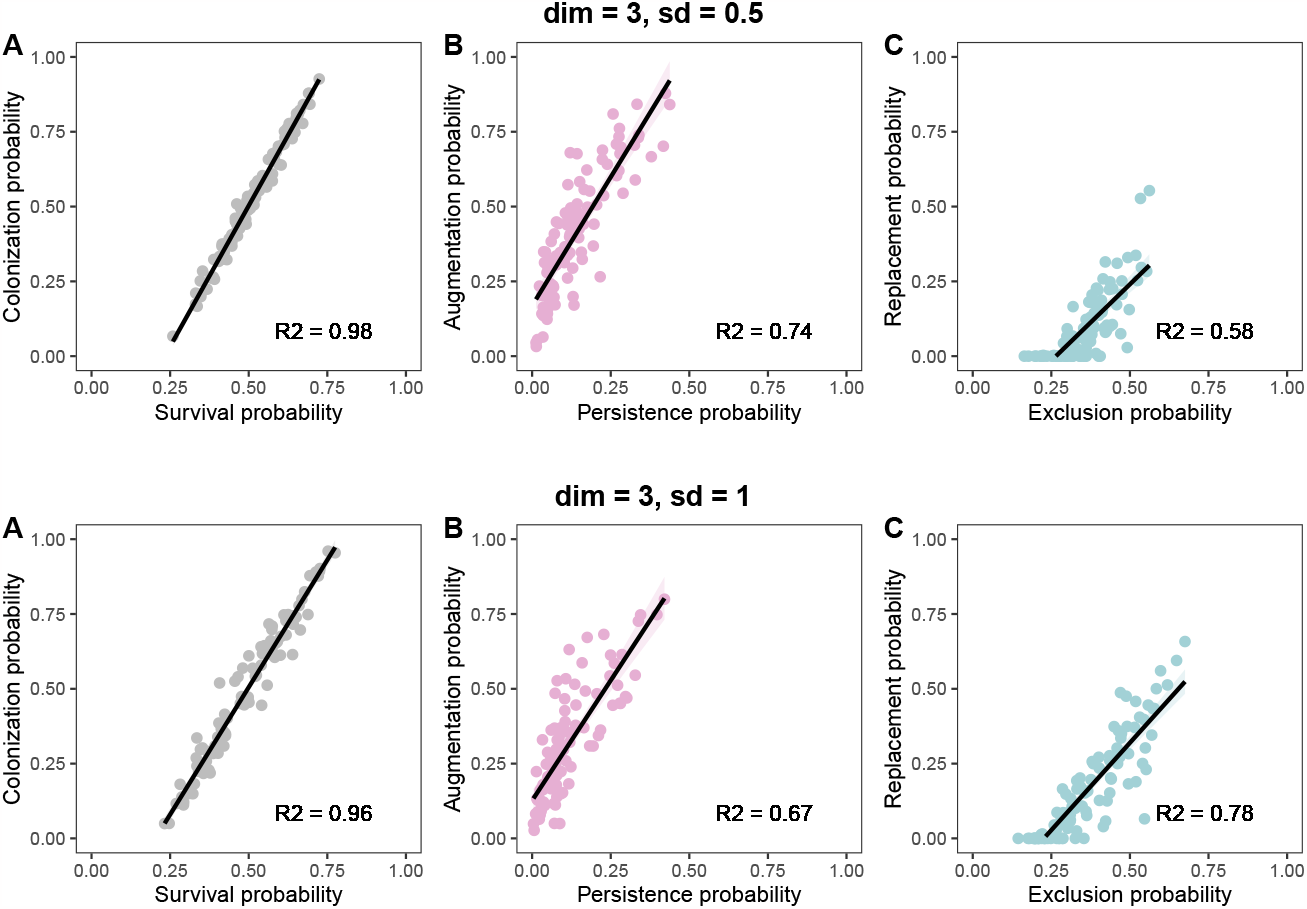
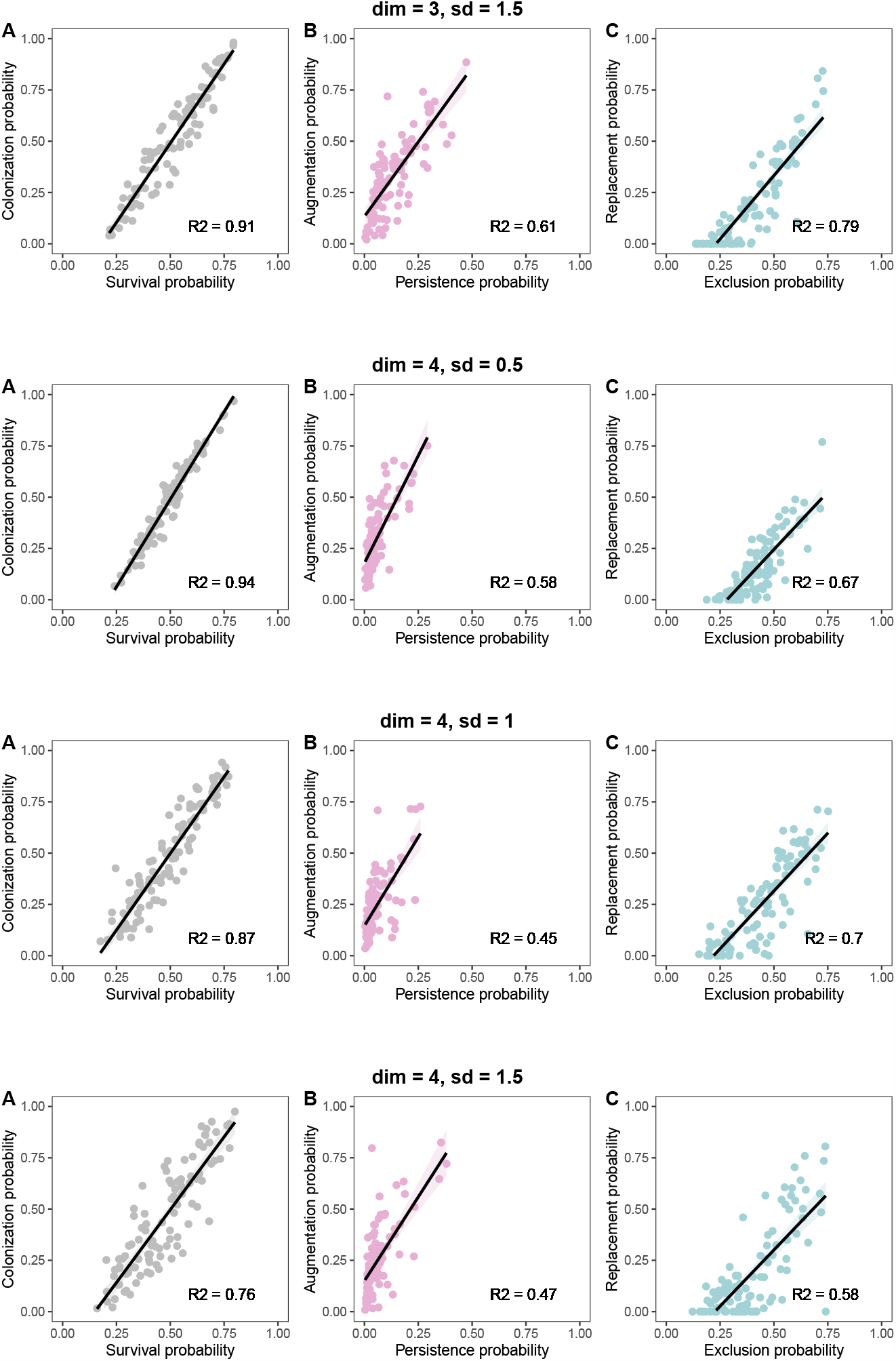
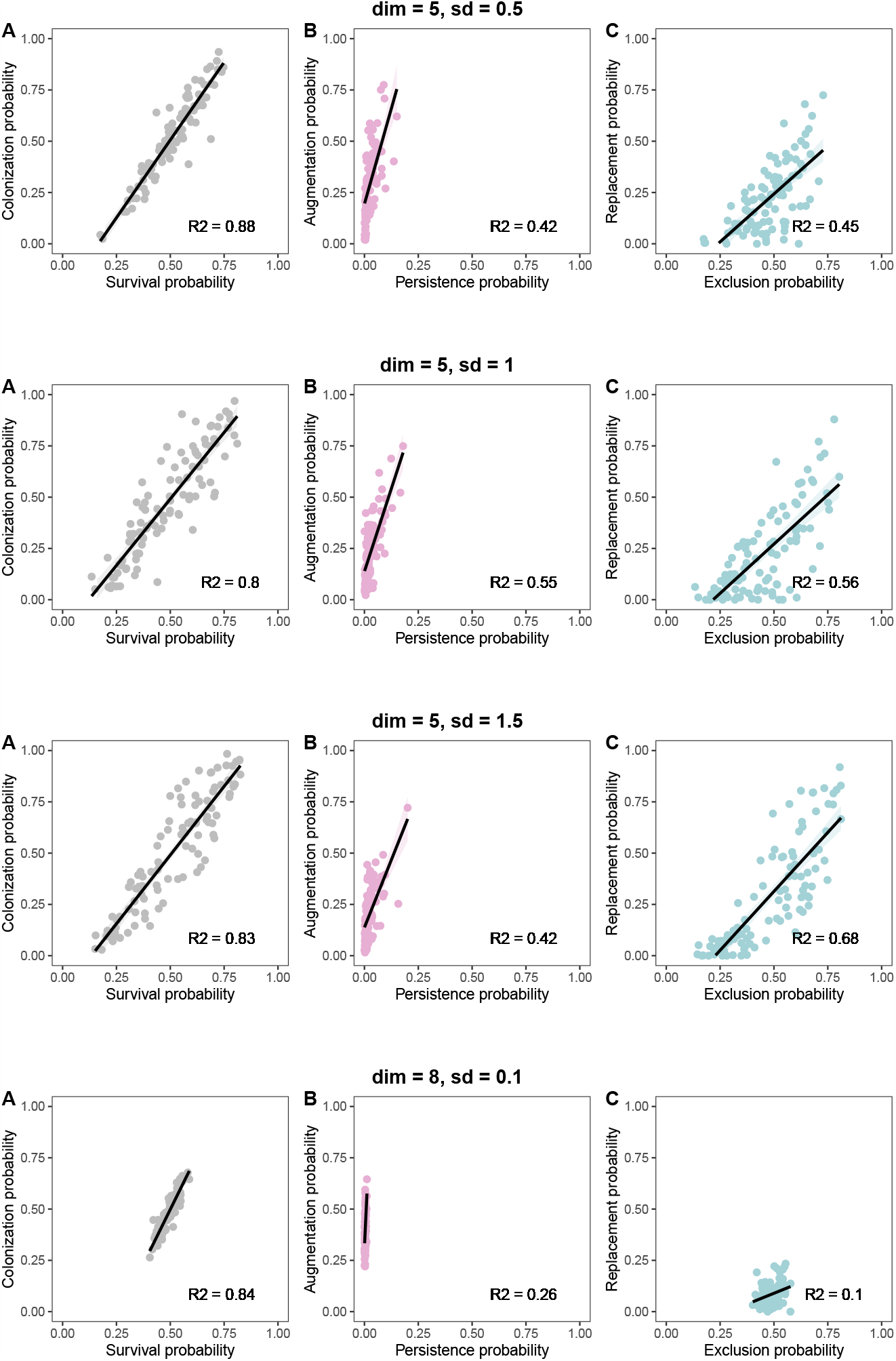
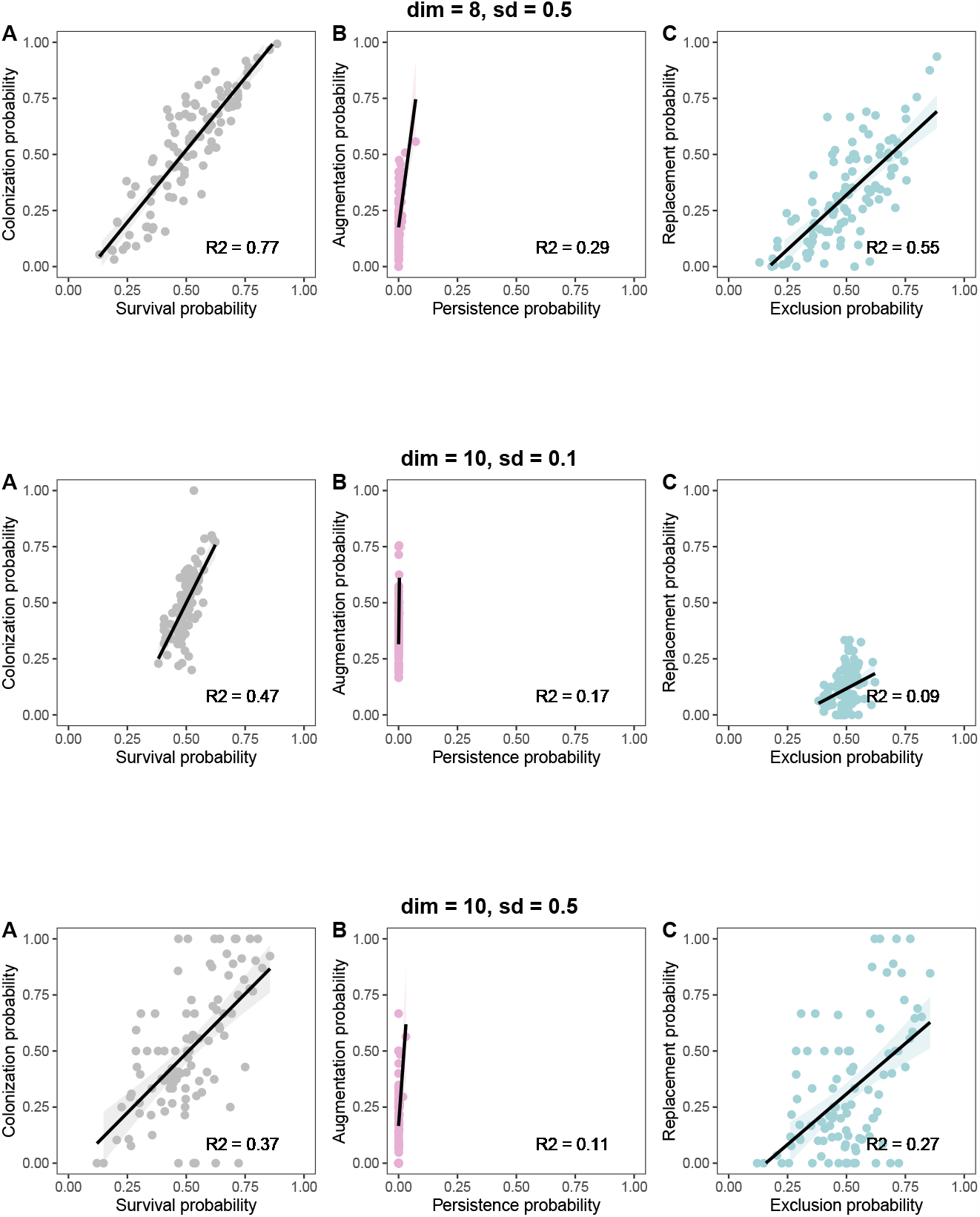
Reproducing Fig. 3 with different numbers of species (dim) and different standard deviations of the interactions (sd). The figures illustrate the numerical results using gLV simulations of three pairs of analogous probabilistic outcomes based on 100 randomly generated communities. The x-axes and y-axes correspond to the outcomes under coexistence and invasion dynamics, respectively. Each point corresponds to a community with the number of species (dim) *∈ {*3, 4, 5, 8, 10*}*. The interaction matrices **A** are randomly generated using a normal distribution with mean *μ* = 0 and standard deviations (sd) *∈ {*0.1, 0.5, 1, 1.5*}*(diagonal entries *a*_*ii*_ =*−* 1). Specific parameter combinations are shown in the caption of each figure. In summary, the numerical results with different combinations of parameters are qualitatively the same as Fig. 3.

## S4 Locally stable communities

Considering three-species locally stable communities, we sampled 50 interaction matrices **A** from a normal distribution with mean *μ* = 0 and standard deviation *σ* = 2 (diagonal entries *a*_*ii*_ = *−*1). Then, for each locally stable community, we sampled 5 *×* 10^3^ vectors of effective growth rates ***θ*** uniformly on a three-dimensional closed unit sphere corresponding to the unknown and changing environments. Also, we set the initial abundance of the invader (focal) species *i* as 10^*−*3^, and draw that of the other two species from a uniform distribution with limits min = 0 and max = 1. Next, to check the surviving species in each community, we simulate the Lotka-Volterra model (Eq. (1)) with total time 200 and step size 10^*−*2^. The extinction threshold is 10^*−*6^. In particular, for invasion dynamics, we perform separate simulations for the resident communities with interaction sub-matrices 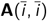, truncated effective growth rates 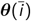, and corresponding initial conditions to guarantee that resident species can coexist in isolation (see the definition of the feasibility domain of resident species in isolation 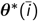 in Eq. (9) and the geometric illustration in Fig. 1A). The simulations are conducted by the Runge-Kutta method. Hence, the probability *F* (*𝒞, 𝒮*) in coexistence dynamics (Eq. (8)) and the probability *I*_*i*_ (*𝒞, 𝒮*) in invasion dynamics (Eq. (9)) of a community 𝒞 within a multispecies community 𝒮 can be approximated by the frequency with which all species in the community 𝒞 coexist together. Consistent with the main text, we use the *R*^2^ to measure the translatability of analogous outcomes between coexistence and invasion dynamics. Fig. S2 illustrates the numerical results of three pairs of analogous probabilistic outcomes. Specifically, Panel A shows the pair of individual survival and invader colonization probabilities (*R*^2^ = 0.94). Panel B shows the pair of community persistence and community augmentation probabilities (*R*^2^ = 0.94). Panel C shows the pair of exclusion and replacement probabilities (*R*^2^ = 0.84).

**Supplementary Figure S2:**
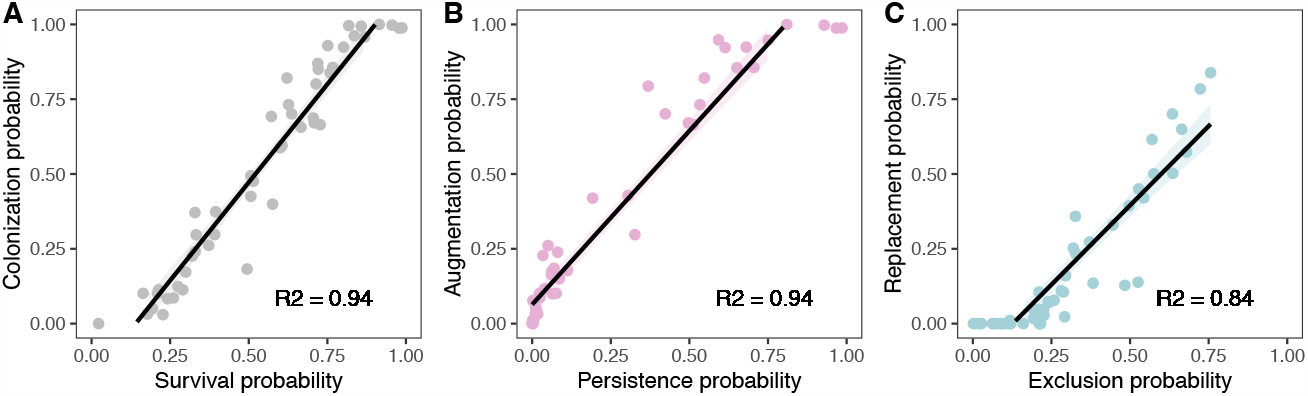
Relationship of analogous outcomes between coexistence and invasion dynamics in locally stable communities. The figure illustrates the numerical results of three pairs of analogous probabilistic outcomes using 50 locally stable three-species communities that are randomly generated. The x-axes and y-axes correspond to the outcomes under coexistence and invasion dynamics, respectively. Each point corresponds to a three-species community, whose interaction matrix **A** is randomly generated using a normal distribution with mean *μ* = 0 and standard deviation *σ* = 2 (diagonal entries *a*_*ii*_ =*−* 1). All probabilities are calculated numerically using an ensemble of 5 *×* 10^3^ effective growth rates ***θ*** sampled uniformly inside the sample space (i.e., a three-dimensional closed unit sphere for coexistence dynamics and the feasibility domain of resident in isolation for invasion dynamics). The results are qualitatively similar to the globally stable cases in Fig. 3 and Supplementary Section S3.

